# Diverse operant control of different motor cortex populations during learning

**DOI:** 10.1101/2021.10.29.466535

**Authors:** Nuria Vendrell-Llopis, Ching Fang, Albert J. Qü, Rui M. Costa, Jose M. Carmena

## Abstract

During motor learning, as well as during neuroprosthetic learning, animals learn to control motor cortex activity in order to generate behavior. Two different population of motor cortex neurons, intra-telencephalic (IT) and pyramidal tract (PT) neurons, convey the resulting cortical signals within and outside the telencephalon. Although a large amount of evidence demonstrates contrasting functional organization among both populations, it is unclear whether the brain can equally learn to control the activity of either class of motor cortex neurons. To answer this question, we used a Calcium imaging based brain-machine interface (CaBMI) and trained different groups of mice to modulate the activity of either IT or PT neurons in order to receive a reward. We found that animals learn to control PT neuron activity faster and better than IT neuron activity. Moreover, our findings show that the advantage of PT neurons is the result of characteristics inherent to this population as well as their local circuitry and cortical depth location. Taken together, our results suggest that motor cortex is optimized to control the activity of pyramidal track neurons, embedded deep in cortex, and relaying motor commands outside of the telencephalon.

## Introduction

To execute a complex natural or neuroprosthetic behavior, animals learn to control the activity of neuronal circuits on motor cortex and the commands sent downstream. Dopamine dependent plasticity between output neurons of those neuronal circuits with its projecting regions seems to be critical for learning (Athalye et al., 2020; Koralek et al., 2012). However, it is still unclear whether there is any difference when learning to control different subpopulations of motor cortex neurons that may have different projecting regions. Understanding the impact that cell-classes have over learning and the different behaviors that they elicit will give insight on how learning may be implemented in the brain.

Motor cortex output is mainly dominated by two cell-classes: intra-telencephalic neurons (IT) which project to the contralateral cortex and bilaterally to the striatum; and extra-telencephalic or pyramidal-tract neurons (PT) which project ipsilaterally to the striatum, to the brainstem and to the spinal cord. Aside from their different projection targets, they also differ in morphology (Wilson, 1987), connectivity (Harris and Shepherd, 2015) and activity (Beloozerova et al., 2003; Cowan and Wilson, 1994). However, there is little information about the extent to which these cell-classes are involved in the learning processes. PT neurons are exceptionally well positioned to generate a cortical output to downstream areas (Egger et al., 2020; Takahashi et al., 2020). Nevertheless, being recruited for such important mission could constrain their flexibility to adapt and accommodate new neuronal patterns. On the contrary, IT neurons are more adaptable (Harris and Shepherd, 2015; Shepherd, 2013), which should be advantageous for efficient learning in neuronal circuits. It is also possible that this division, IT vs PT, could be less relevant to neural control and manipulation than other neuronal characteristics such as location, activity or circuit dynamics of neighboring neurons. Understanding what drives the adaptive mechanisms of control over cortical neurons is not only relevant for clarifying the role of IT and PT neurons during learning, but it can also enlighten the different roles of cortical cell-classes in disease (Shepherd, 2013) and motor function (Li et al., 2015; Reiner, 2010).

To address these questions, we took advantage of an established calcium imaging brain-machine interface (CaBMI) paradigm (Clancy et al., 2014) while utilizing viral tracing to probe the functional properties of different cortical cell-classes across multiple cortical layers in behaving mice. A CaBMI paradigm relies on an operant learning task where mice volitionally control their neuronal activity in order to obtain a reward. Specifically, mice learned to control either IT or PT neurons allowing us to address any difference in the ability of the brain to learn with either neuronal population. Finally, with machine learning and game theory approaches we dissected all possible influences on learning and showed the relevance of the inherent characteristics of each cell-class and their local circuitry and their positive influence on learning.

## Results

To distinguish IT neurons from PT neurons, we engineered cell-class specific expression of a red fluorescence marker in two groups of tetO-GCaMP6s/-Camk2a-tTA transgenic mice. One group (n=9) was injected with AAVrg-CAG-tdTomato in the contralateral motor cortex to label IT neurons and the other group (n=8) in the ipsilateral pons to label PT neurons (Fig.1.A-E). We trained both groups of mice to control a CaBMI (see methods) while simultaneously recording in four different planes the activity of both red-labeled and unlabeled neurons (Supp.Fig.1.A-C). Because the recordings span a large part of the cortical column (400µm, Supp.Fig.1.D), unlabeled neurons may contain excitatory neurons from both cell-classes and possibly inhibitory neurons (Nathanson et al., 2009; Watakabe et al., 2015). Each session, a different pair of red-labeled neurons were arbitrarily selected and assigned to neural ensembles. By using only PT or only IT neurons as the neurons directly controlling the CaBMI (direct neurons), we studied the differences in learning between these genetically different subpopulations of cortical neurons as well as the neural dynamics that surrounded them.

**Figure 1:**
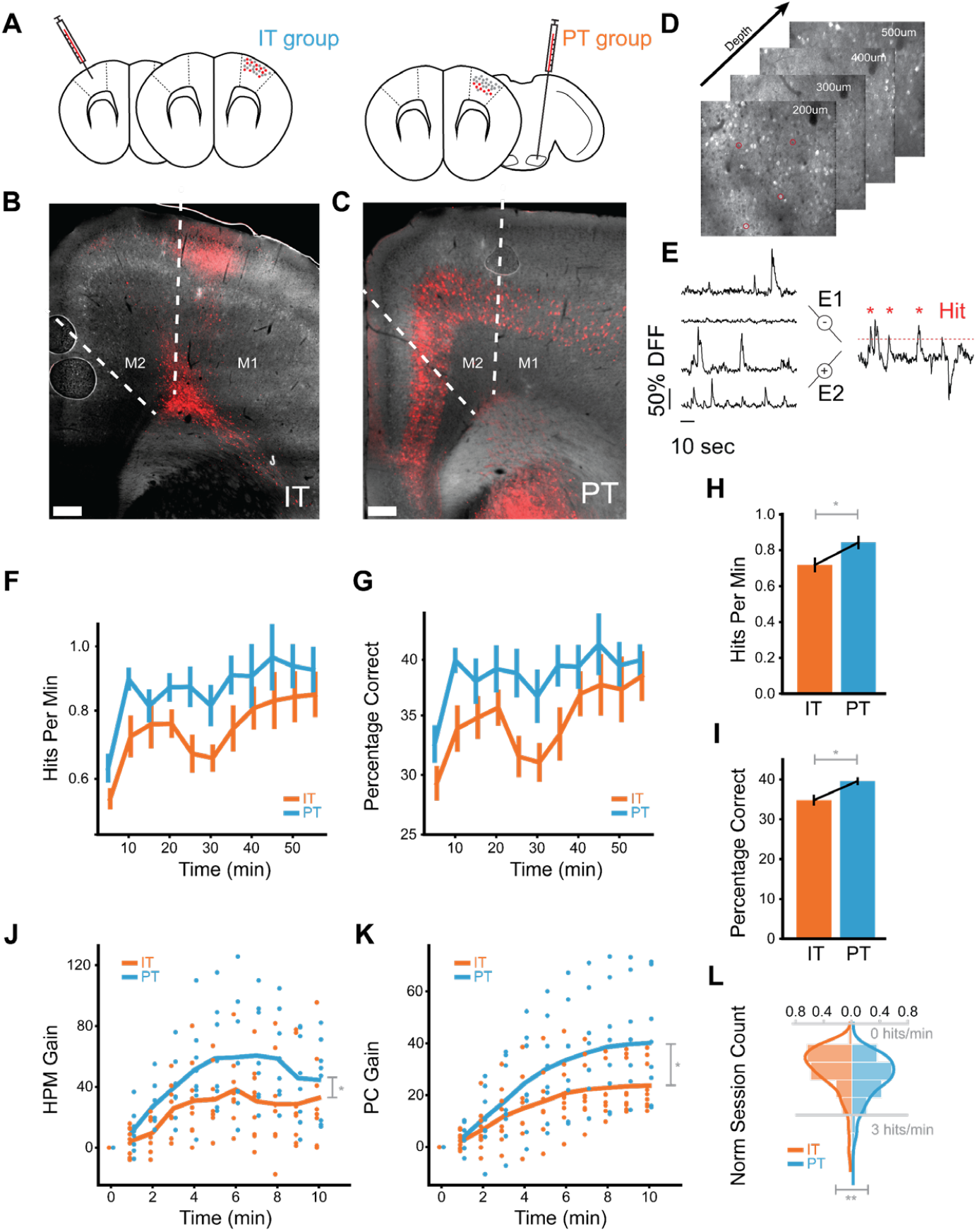
Cell-class specific CaBMI shows differences on learning. **A**) AAVrg-CAG-tdTomato retroviruses injection regions for IT (left), contralateral motor cortex, and PT (right), ipsilateral pons. (**B-C**) Coronal sections for the IT (**B**) and PT (**C**) animals. Neurons with viral expressions were located in L2/3 and L5-6 of motor cortex (M1-M2) for IT group and in L5-6 across cortex for PT. **D**) Typical depth planes for calcium recordings. Marked neurons were randomly picked as direct neurons. **E**) Schematics for the calcium decoding algorithm. Neural cursor is calculated as the difference between sum dF/Fs of two groups. Red dashed line denotes one instantiation of the reward threshold simulated from baseline. **F**) HPM, the number of hits within a one-minute window, calculated for each session across IT experiments (orange) and PT experiments (blue). Values are binned in five-minute windows. **G**) As in (**F**), but for PC, the percentage of correct trials within the same one-minute window. Because trial lengths can be variable, PC represents the reward rate normalized by the number of trials in each time window. **H**) HPM values across all time windows of a session (*: p < 0.05 with Mann-Whitney U test). HPM chance level = 0.46 ±0.04. **I**) As in (**H**), but for PC. PC chance level 0.25±0.02 **J**) The relative gain in HPM during the first 10 minutes of a session from the beginning of the session. Specifically, we calculated the difference in HPM from the first minute of the experiment, then normalized this measure for each session by dividing by the mean HPM. This normalization allowed us to compare performance increases across sessions with different starting reward rates. **K**) As in (**J**), but for PC. **L**) Distribution of maximum HPM obtained per session. This measure of performance is less affected by periods of low motivation or attention, which can modulate learning curves. (**: p < 0.005 with one-way Anova)

Each session we changed the direct neurons to test learning capabilities in as many neurons as possible. For that reason, our results can only be compared to initial sessions of other CaBMI experiments (Athalye et al., 2018; Clancy et al., 2014; Hira et al., 2014; Mitani et al., 2018; Prsa et al., 2017). By using naive neurons, we investigated the role of IT and PT neurons during the acquisition of a learned behavior which may entail different processes and circuits than the refinement of that behavior (Athalye et al., 2020). The role of different cortical neurons in later stages of learning including consolidation and refinement should be addressed in further experiments.

### Animals learn to control pyramidal tract neurons better and faster than intra-telencephalic neurons during CaBMI learning

We first investigated whether our choice of IT or PT neurons as direct neurons affected the animal’s learning ability. We quantified learning through two measures: hits-per-minute (HPM) and percentage-correct (PC). Hits-per-minute quantifies reward rate over time while percentage-correct is the reward rate normalized by the number of trials. Both groups showed an increase from chance level (HPM: 0.46±0.04, PC: 0.25±0.02) in both hits-per-minute and percentage-correct throughout the experiment (Fig.1.F-G). However, the PT group (n=125 sessions) achieved greater reward rates than the IT group (n=162 sessions) across equivalent time windows of a session (Fig.1.F-G) and across the whole session (Fig.1. H-I).

To address if differences in starting reward rate explained the learning differences between the IT and PT group, we obtained the gain of each learning measure over the first 10 minutes. We found that the PT group reached a higher performance than the IT group and did so faster (Fig.1.J-K). Similarly, we compared the best performance of each group to address possible effects of motivation loss. We observed that the PT group tended to have significantly higher maximum performance (p<0.005, Fig.1.L).

Taken together, our results demonstrate that, although both groups can learn the task, the PT group consistently outperforms the IT group across a variety of learning measures. Strikingly, the PT group achieves a higher reward rate than the IT group in less time. Much of this performance difference occurs in the first few minutes of an experiment (Fig.1.J-K). A possible explanation is that the network involved in this task learns to modulate the activity of PT neurons and/or re-enter the neuronal patterns that granted reward, more effectively and quickly. If so, known characteristic differences between IT and PT neurons, such as connectivity or cortical depth (Reiner, 2010), may be relevant. However, other factors besides neuron properties can also contribute to these observed differences. Experimental confounds, such as imaging quality, may explain these differences. In the rest of the paper, we tackle the extent to which a diverse range of factors could explain the higher learning performance of the PT group.

### Evaluating experimental and neural features that influence learning

To control for possible experimental confounds and understand the impact of cell-class and other neural features on learning, we implemented a feature attribution framework. We used a gradient-boosted decision tree, XGBoost (Chen and Guestrin, 2016), as a model to predict the percentage of correct trials (henceforth percentage-correct), our measure of learning. Then, we used SHapley Additive exPlanations (SHAP) to explain how each feature contributed to the model’s prediction each session (Supp.Fig.2). Thus, positive SHAP values represent that a feature pushed the predicted value of percentage-correct higher, suggesting a positive influence on learning. On the contrary, negative SHAP values represent that a feature reduced the predicted value of percentage-correct, suggesting a negative influence on learning (Fig.2.A).

**Figure 2:**
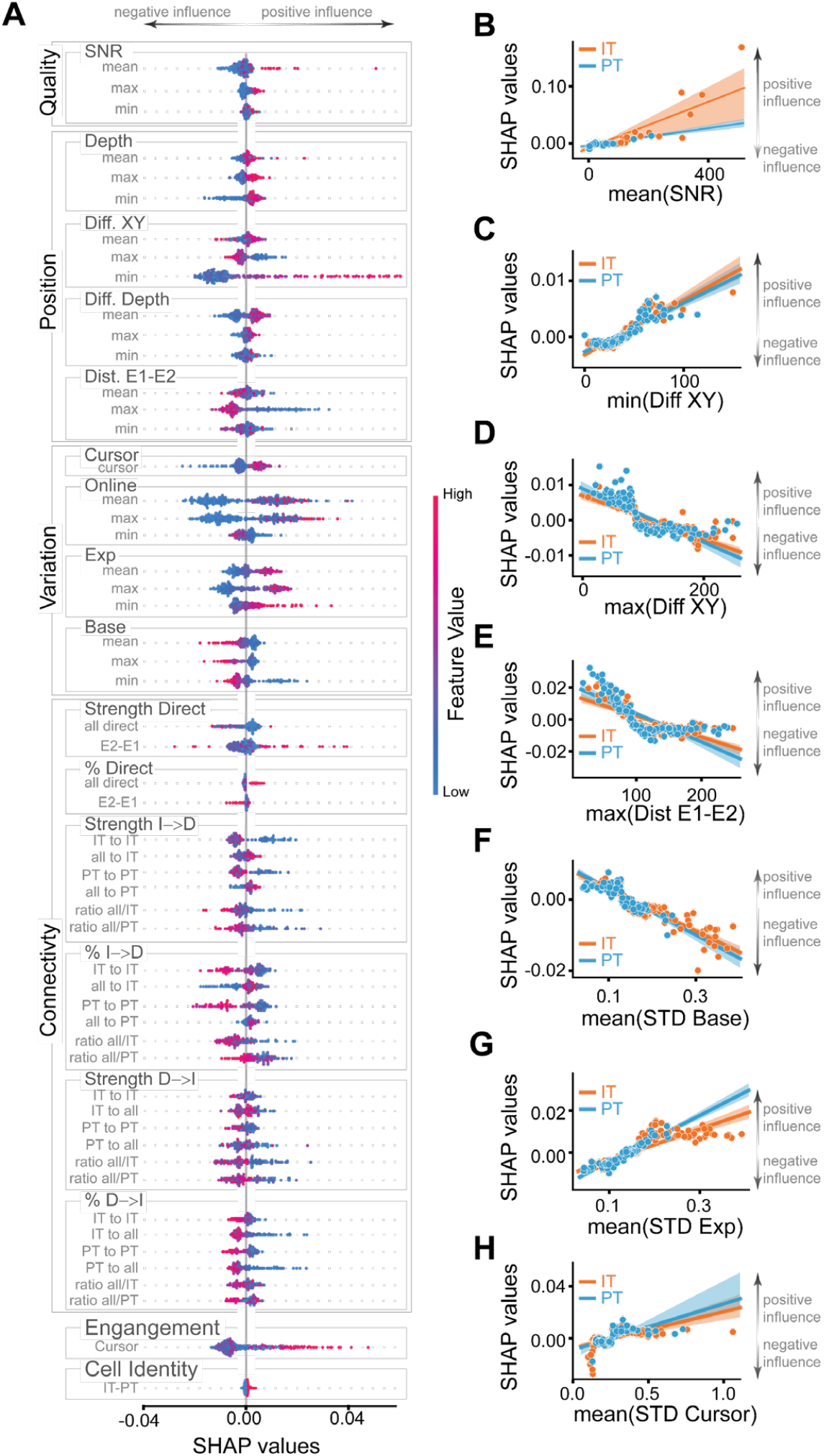
Dissecting the influence of features on learning outcome. **A)** SHAP values for all the features used in the XGBoost models (see Supp.Fig.2). Positive SHAP values indicate a positive effect on the PC measure and a better learning outcome. Negative SHAP values indicate a negative effect on learning. Features are divided into 6 groups: Quality, Position, Variation, Connectivity, Engagement and Cell Identity for clarity. Quality features are comprised of the signal-to-noise ratio. Position features include depth (Depth) of direct neurons, the distance between direct neurons on a plane (Diff. XY), their difference in depth (Diff. Depth), and the distance between neurons of the ensemble E1 and the ensemble E2 (Dist. E1-E2). Variation features include STD of the cursor (Cursor) based on neuronal activity E2-E1, STD of the online calcium signals that operated the CaBMI (Online), and also the post-processed calcium signals of direct neurons during baseline (Base) or during the whole experiment (Exp). Measures that were calculated for each direct neuron were included as features with their minimum (min), mean, or maximum (max) value. Color (red-to-blue) represent the value of the feature. Each dot represents 1 session. Sessions from IT and PT groups were included. **B-I)** Linear regression between SHAP values and the value of mean SNR (**B**); minimum distance (**C**) and maximum distance (**D**) in the XY plane between 2 direct neurons; maximum distance between neurons belonging to different ensembles (**E**). Activity of direct neurons during baseline (**F**) or whole experiment (**G**). **H)** Mean of the STD of the neural cursor. Each dot is a session. Shaded area is the confidence interval. Orange dots are sessions from the IT group whereas blue dots are sessions of the PT group.

Untangling the true effect each feature had on the animal’s learning each session is a complicated endeavor. For example, depth, signal quality and cell identity are highly correlated but may have opposite effects on learning. By using a highly accurate model and a reliable explainer we offer a reasonable approximation to the immeasurable independent contributions of each feature to learning.

### Experimental-dependent features and cell-class activity variability do not explain learning differences

We first investigated if experimental-dependent features (features that were solely affected by experimental limitations or experimenter bias) affected learning. These features captured signal quality and distance between direct neurons (excluding cortical depth which is dependent on cell-class). SHAP values were highly correlated with signal-to-noise ratio (SNR), indicating that cleaner signals resulted in better animal performance (Fig.2.B). Thus the generally lower signal-to-noise ratio of PT neurons (due to brain-scatter imaging limitations in deep tissue) hindered the predicted percentage-correct (Supp.Fig.3.A-B). Additionally, the distances between direct IT or PT neurons (Fig.2.C-E) affected the value of the predicted percentage-correct similarly in both groups. These results indicate that none of the experimental features that were independent of cell-class were responsible for the improved performance of the PT group.

Distinct activity characteristics have been identified in IT and PT neurons (Dembrow et al., 2010). To investigate if these differences could influence learning, we added features based on the standard deviation (STD) of neuronal activity to the model. Highly active direct neurons during baseline negatively affected the value of predicted percentage correct (Fig.2.F). Contrarily, highly active direct neurons during the experiment increased the predicted percentage correct (Fig.2.G). These apparently contradicting results are consistent with CaBMI benefiting from silent neurons during baseline becoming more active during the task. The generally tonic firing characteristics of PT neurons (Dembrow et al., 2010) resulted in reduced changes of fluorescence in calcium imaging and therefore low variability (Supp.Fig.3.C-E). These had contradicting effects on the predicted percentage-correct (Supp.Fig.3.D-F). In addition, there was no cell-class difference due to the variability of the CaBMI cursor (Fig.2.H). These findings indicate inconclusive effects of the different activity characteristics of PT and IT neurons over learning.

### Local connectivity and position of PT neurons accounts for the differences in learning

Learning modulates the activity of indirect neurons (Ganguly et al., 2011; Zippi et al., 2021), neurons of the local circuitry recorded during the experiments but not in direct control of the CaBMI. However, the influence that local circuitry may pose on learning has not yet been investigated. To address this, we studied how two different measures, connectivity to/from direct neurons and task engagement of indirect neurons, would influence learning.

To begin, we used Granger causality as a measure of effective connectivity. We determined the percentage of neurons that were connected as well as the strength of those connections, during the baseline period (see methods). It is important to note that this method cannot capture the effects of neurons not recorded and/or fast recurrent networks due to our recording framerate (10Hz). Higher effective connectivity (both in the number of pairs and the strength of those connections) from direct neurons to indirect neurons, regardless of cell-class, lowered the predicted percentage-correct. On the contrary, higher connectivity (also in the number of pairs and strength) from unlabeled indirect neurons to both IT and PT direct neurons increased the predicted percentage-correct (Fig.2.A), suggesting better performance for neurons receiving higher, in both number of connections and strength of those connections, indirect input.

Lastly, to evaluate how engaged were the indirect neurons with the task, we calculated how well these indirect neurons could predict the neural cursor controlling the CaBMI (see methods). Besides some fluctuations, better cursor prediction by indirect neurons was tightly linked with higher predicted percentage-correct (Fig.2.A), indicating a positive influence in learning.

These findings suggest that generally, a strong support from local circuitry positively impacts learning. However, our results are limited to the acquisition of a learned behavior. Previous research have shown that during late learning, task modulation of indirect neurons decreases (Ganguly et al., 2011; Zippi et al., 2021) and there is less functional connectivity from indirect neurons to direct neurons than vice versa (So et al., 2012). It is possible that local circuitry may be more relevant during early learning facilitating the re-entrance of the neuronal patterns that grant reward. On the contrary, the effects of refinement may entail different neural processes, including the pruning of indirect neurons deemed impractical.

Interestingly, both measures quantifying the involvement of indirect neurons (effective connectivity and task engagement) increased the predicted percentage-correct for the PT group but decreased it for the IT group (Fig.3.A-B). This may imply that deep local circuitry surrounding PT neurons more effectively supports learning than circuitry in the upper cortical layers. Is this also true for IT neurons? To investigate this, we examined if choosing IT direct neurons from deeper planes affected learning outcomes. We found that sessions with IT direct neurons in deeper cortical layers increased the predicted percentage-correct (Fig.3.C). Additionally, depth was highly correlated with indirect-to-direct connectivity (Fig.3.D, Supp.Fig.3.I) which our results indicate facilitates learning. Taken together, these findings suggest that direct IT neurons located in the vicinity of PT neurons learned more effectively than their counterparts in upper layers. However, it is possible that this learning difference arises not from a supportive deep local circuitry but from the postsynaptic circuit. IT neurons projecting into striatum are oftentimes found in deeper layers than IT neurons that only project into other cortical regions (Shepherd, 2013). Thus, the behavioral outcome may be largely influenced by synaptic proximity to the next stop of the cortico-basal-thalamo-cortical loop that governs learning.

**Figure 3:**
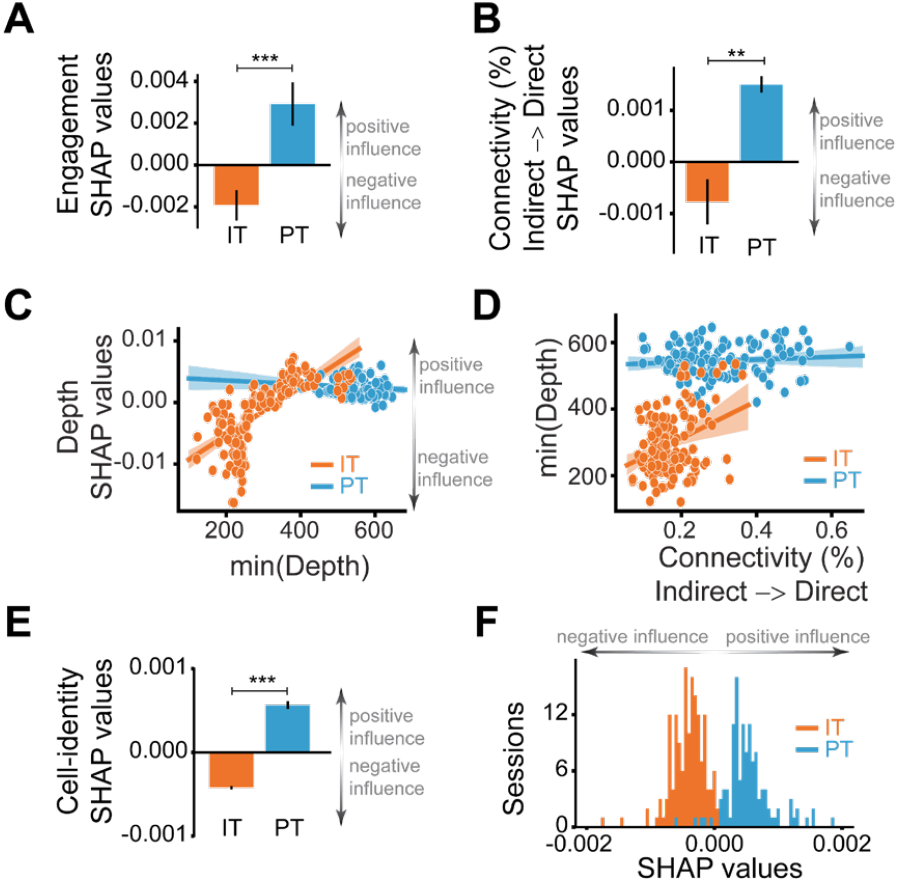
Inherent characteristics, connectivity and location of PT neurons lead to better performance. SHAP values for circuit-related features: (**A**) engagement of indirect neurons and (**B**) effective connectivity from indirect to direct neurons in sessions of the IT (orange) or PT (blue) groups. **C**) SHAP values depending on the minimum depth of all direct neurons. **D**) Linear regression between the minimum depth of all direct neurons and the effective connectivity from indirect to direct neurons. Each dot is a session of the IT (orange) or the PT (blue) groups. Line is the linear regression and shaded area its confidence interval. **E**) SHAP values for the cell-identity feature and their distribution (**F**) with IT group sessions in orange and PT group sessions in blue. Black lines in bar graphs represent SEM. (** : p<0.005, ***: p<0.0005 with independent t-Test).

We aimed to investigate if location is the only relevant feature to determine learning. However, we could not measure all possible different characteristics of IT and PT neurons (such as input from other cortical areas, thalamic input, spike burstiness, etc.). Instead, we added one more feature that encoded the identity of the neurons chosen for CaBMI, hence accounting for the remainder of other inherent cell-class characteristics. Strikingly, the difference between both groups was very consistent. Selecting PT neurons for CaBMI generally increased the predicted percentage correct, indicating a positive influence on task performance whereas selecting IT neurons had an entirely negative effect on the predicted percentage-correct (Fig.3.E-F). This effect indicates that inherent characteristics of PT neurons were consistently advantageous for successful learning.

In summary, by using a method that measures the impact of different cell-type features on the learning of an operant CaBMI task, our work provides insights into the understanding of the factors that contributed to operant control of cortical activity. Our results demonstrate that animals learned to control PT neurons faster and more effectively than IT neurons and that this effect cannot be attributed to any experimental confounds. Instead, our results suggest that the brain is more effective at manipulating and controlling output neurons that project from cortex to regions outside the telencephalon, and that this results from connectivity and position in the cortex.

## Acknowledgments

This work was supported by the NIH U19 grant to RMC and JMC, and the Army Research Office Award W911NF-16-1-0453 to JMC.

## Author contributions

NVL, RMC, and JMC designed the study. NVL and CF performed the experiments. NVL, CF, and AJQ analyzed the data. JMC and RMC lead the study. NVL, CF, and AJQ wrote the manuscript with significant contributions from RMC and JMC.

## Methods

### Animals & Surgery

All experiments were performed in compliance with the regulations of the Animal Care and Use Committees at the University of California, Berkeley and according to NIH guidelines. Mice were housed with a 12-h dark, 12-h light cycle. Two groups of tetO-GCaMP6s/-Camk2a-tTA mice (original strains from The Jackson Laboratory, Bar Harbor, Maine: jax-024742, jax-007004) were injected with a retrograde virus in the contralateral motor cortex (n=9) or in the ipsilateral pons (n=8) in order to label intratelencephalic (IT) neurons and pyramidal tract (PT) neurons, respectively. Although cortico-thalamic neurons would also be relevant to this study, their location deep in the brain, made them impossible to study under the current state of the art of two-photon microscopy.

Prior to surgery, tools and materials were sterilized by autoclaving or gas sterilization. Mice were initially anesthetized by placing them briefly (2-3 mins) in a box containing 3 - 4% isoflurane and were then kept at 1-2% isoflurane in a nose cone respirator connected to a precision vaporizer. The animal was secured into a stereotaxic frame (Kopf instruments, Tujunga, CA) and kept warm (37.5 ± 1 °C). A single incision was made along the midline of the skull in the rostro-caudal direction and the skull was cleaned. Using a rotary micromotor drill (Foredom, Bethel, CT) equipped with a 0.5mm carbon burr (Fine Science Tools, Foster City, CA), a small burr hole was made over the contralateral motor cortex (1.4 mm rostral, 1.3 mm lateral to Bregma) or the ipsilateral pons (4.26 mm posterior, −0.6 mm lateral to Bregma), and 400 nl of AAVrg-CAG-tdTomato (Addgene Watertown, MA, viral prep # 59462-AAVrg) was injected 300um (for IT) or 4.6mm (for PT) below the pia. The tracer was delivered using a pulled glass pipette (tip diameter = 40– 60 μm) at a rate of 50 nl min with a Nanojet 3 (Drummond Scientific Company, Broomall, PA). The pipette was left in the brain for 15 min after completion of the injection to prevent backflow. After removal of the pipette, the burr hole was covered with Metabond dental cement (Parkell Edgewood, NY, S396-S398-S371). A 3-mm craniotomy was opened over motor cortex (coordinates of the center relative to Bregma ML-1.5, AP 1.3). Two sterile glass coverslips (3-5mm, #1 thickness) were glued concentrically to each other (Norland Optical Adhesive, Cranbury, NJ, NOA71) and positioned over the skull so the 3mm coverslip would fit in the craniotomy. Metabond was applied to create a thin seal between the skull and the sides of the cranial window, and a steel headplate was affixed posterior to the coverslips. We allowed 3-4 weeks for recovery and for the expression of tdTomato, before starting the behavioral experiments. Animals with injections in the contralateral motor cortex had IT neurons labeled with tdTomato (IT group), whereas animals injected in the pons had PT neurons labeled with tdTomato (PT group).

### Two-photon imaging

Recordings of calcium imaging were performed with a Bruker Ultima Investigator (Bruker, Millerica, MA) using a Chameleon Ultra II Ti:Sapphire mode-locked laser (Coherent, Santa Clara, CA) tuned to 920 nm. Photons were collected with two GaAsP PMTs for different channels using an Olympus objective (XLUMPLFLN 20XW). Animals were head-fixed over a styrofoam ball (JetBall, PhenoSys, Berlin, Germany) that allowed them to run freely under the two-photon microscope. A piezo controller (400um travel, nPoint, Middleton, WI) allowed the sawtooth recording of 4 different planes with 100um separation for a full sweep of the cortical column. The power of the laser was set so that high quality images of the planes with direct neurons could be achieved without damaging shallow planes. Different imaging fields were used every day. In a given session, the imaged planes spanned 400 microns in depth. These planes were centered ∼350-550 microns below pia depending on the session. Frames of 256 × 256 pixels (∼290 × 290 μm) were collected at 9.7 Hz using ScanImage software (Vidriotech, Ashburn, VA). Motion drifts (if any) were corrected online by the software and/or manual control. Motion artifacts resulted in poor task performance (since both ensembles moved accordingly) and the mice seemed to remain more still during late learning sessions. Additionally, we added the quality of the recorded calcium signals of the direct neurons (measured as SNR) as features of the XGBoost-SHAP models. SNR was positively correlated with SHAP values, indicating that animals performed better in sessions with higher signal quality.

### Behavioral task and online processing

This behavioral task has been described previously in electrophysiology (Koralek et al., 2013, 2012; Neely et al., 2018) and calcium imaging(Clancy et al., 2014).. Activities of two pairs of M1 neurons were summed within ensemble (∑E1 –∑E2) and entered into a decoder that mapped neuronal activity to an auditory signal (range 2-18kHz). Head-fixed mice could increase the frequency of the auditory cursor by increasing the activity in the first ensemble (E1) and decreasing the activity in the other ensemble (E2). Mice could instead decrease the frequency of the auditory cursor by decreasing the activity of E1 and increasing the activity of E2. Mice received reward (20% sucrose) if they decreased the cursor frequency under a predefined target. To set the target cursor frequency, neuronal activity was recorded during a baseline period of 15 minutes. Each day the target was set such that mice would have received reward in 30% of trials in a hypothetical simulation with the recording from the baseline period. The auditory signal was proved to the animals as feedback of their performance.

In each group of animals (IT or PT), only tdTomato labeled neurons were used to control the auditory cursor. To study within-session learning, different IT or PT neurons (respectively) were selected each day. The animals had 30 seconds to reach the target and achieving a “hit”. Otherwise, the trial would be considered a “miss”. With a successful trial, sucrose reward was given to the animal. After a 3 second pause, the auditory cursor was required to return to a baseline value in order to start the next trial. If the animal did not hit the target in the allowed time, white noise was indicative of fail and the mice were given a 10 second timeout before a new trial started.

Neuron segmentation for online processing was obtained by a template matching function(Ohki et al., 2005). Fluorescence change of each of the identified neurons, defined as (Ft-F_0_)/F_0_ or dF/F was obtained online as a measure of neuronal activity. F_0_ was calculated dynamically to avoid bleaching effects without compromising processing time as F_0_ = (n-1) * F_0_ /n + F_t-1_ where n was the number of frames acquired. For online processing, the F_t_ value of each M1 neuron was averaged over the last second before calculating dF/F, to provide robustness against motion artifacts.

### Image preprocessing

Each of the 4 imaging planes was separated into a block and independently analyzed with CaImAn (Giovannucci et al., 2019) to obtain the activity of each neuron during the recording. Direct neurons selected during the online experiment were matched with CaImAn-identified neurons by activity and space correlation. If a direct neuron could not be matched to a CaImAn-identified neuron (i.e., the activity or position was too different from online ensemble), the neuron was assumed to have had a low SNR and would be removed from the ensemble, reducing the post-hoc ensemble to one neuron. If both neurons were discarded from an experimental session, the session was not used for analysis. Because the positions given by CaImAn are dependent on the whole spatial filter of the neuron and not the soma, new positions were obtained by filtering the image to locate the center of each neuron soma. To identify which CaImAn-identified neurons corresponded with tdTomato labeled neurons, the positions obtained by the template matching function(Ohki et al., 2005) (over the red channel image) were matched to the positions of CaImAn-identified neurons if the Euclidean distance between both centers were less than 4 pixels.

### Data analysis

Analysis programs were custom-written in Python using a variety of packages. The analysis pipeline was consistent across all animals. Code will be made available upon request.

### Signal to noise ratio

Online recordings of direct neurons saved by ScanImage during experiments were used to calculate the online signal to noise ratio (SNR), where 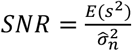. Since ScanImage may drop frames of data during online collection to achieve the desired image rate, we filled the missing frames using linear interpolation and nearest neighbor extrapolation for post-processing. To disentangle noise power from signal power, we averaged over the high frequency ranges (*f*/4, *f*/2with *f* as frame rate) of the raw trace’s power spectral density (Pnevmatikakis et al., 2016). To validate the method’s efficacy, we simulated noisy calcium traces with different noise and bleaching conditions and found that this method, compared to other SNR estimations, better minimizes the L2 norm of the error from predicting ground-truth SNRs.

### Cursor engagement analysis

The cursor engagement value for an experiment is a measure of how well the activity of indirect neurons can predict the auditory cursor. We used L1-regularized linear regression to predict the cursor with the fluorescence of indirect neurons at each frame. Specifically, for each experiment, we collect the ΔF/F values of the *S* indirect neurons over the *T* frames of the experiment into a matrix *X*(*S* ×*T*). The auditory cursor over the *T* frames is collected into a *T* -length vector 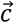. Thus, for each experiment we obtained *T* samples of data with *S* features. For some frame *t*{1, …, *T*},the goal was then to predict 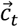 with *X*_·{1:*t*−1}_. We first split the *T* samples 80/20 into a training set and a testing set. The training set was used to train the model and select hyperparameters with 5-fold cross validation.

The model’s performance was then evaluated on the testing set. The quality of the testing set prediction was quantified by the *R*^2^ coefficient of determination value. The best possible *R*^2^ value is 1. A constant model that always predicts the expected value of the cursor would have *R*^2^ =0. A model that does worse than this constant model would have *R*^2^ <0. Since the r^2 value can become arbitrarily negative and since models with *R*^2^ ≤0 were ineffective in predicting the cursor, the cursor engagement for an experiment was calculated as max{0,*R*^2^}. This allowed the cursor engagement value to lie in a predefined range.

### Granger Causality

We used Granger causality to estimate the bi-directional effective connectivity between each pair of tdTomato-labeled neurons and between each direct neuron and indirect neurons. Granger causality models time series as autoregressive series. A trace x is said to be “Granger causal” to y if, given the following two formulations:

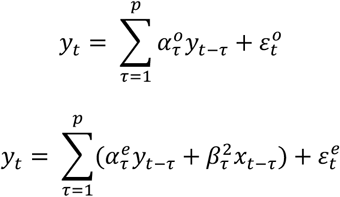

the Granger causality value 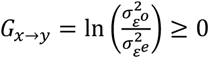 (with equality achieved when *x*_1:t_⊥⊥ *y*_1:t_).

We selected an autoregressive model of order *p* = 2 based an average case of order selection by minimizing Bayesian information criterion.

Then, for each directed pair, we used chi-squared tests on sum of squared residuals (SSR) to determine the statistical significance of the directed influence. Only estimated effective connectivity values for neuron pairs with p-value less than 0.05 were kept as raw features.

To determine the effectiveness of Granger causality inference algorithm in reconstructing effective connectivity for calcium data, we performed the following two validations. First, we simulated a series of excitatory neural networks with Integrate-And-Fire neurons with connectivity determined by an Erdos-Renyi graph *g*(*n, p*) with different n, p parameters. We converted the simulated spike data into calcium data with the Leogang model (Stetter et al., 2012). We then processed the simulated data with the granger causality and obtained the Area-Under-Curve for the Receiver Operating Characteristics graph over chance level. Second, we validated Granger causality’s efficacy on calcium data by comparing the connectivity values among neuron pairs to values among shuffled pairs, To generate realistic random activities with comparable statistics, we obtained shuffled calcium data by re-convolving shuffled deconvolved spikes. As a control for artifacts introduced by deconvolution, we also re-convolved all unshuffled spike data and calculated their inferred connectivity to compare against the shuffled version.

### XGboost/SHAP

SHAP values(Lundberg and Lee, 2017), were obtained for XGBoost (eXtreme Gradient Boosting) models (Chen and Guestrin, 2016) with a TreeSHAP (Lundberg et al., 2020) for each of the features and experimental sessions in the following manner. 10000 models were trained on 80% of the experimental sessions and tested on the remaining 20% with XGBoost using random sampling with replacement. XGBoost models regressing percentage-correct values (average mean square error = 0.026, representing less than 7% of the average percentage-correct) outperformed XGBoost models regressing hits-per-minute values (average mean square error =0.27 ∼ 35%). Thus, we selected percentage-correct as the learning measure to regress and all following analysis was done only for percentage-correct models. Only models with high accuracy and low variance were chosen for further analysis (see below). Parameters for the XGBoost models were chosen to maximize the accuracy of the model although varying them only affected accuracy slightly (learning _rate=0.1, repetitions=100, Bootstrap repetitions=1000).

SHAP values were obtained with TreeSHAP for the test data only. We used “tree_path_dependent” as feature perturbation to remain true to data (Chen et al., 2020). Because we obtained 10000 different models within specifications, each experimental session was part of the test data more than once, resulting in multiple SHAP values for each experimental session and feature. Each session was used in a model an average of 2010 times. However, 3 sessions with high performance (PC = 0.9151, 0.9202 and 0.8519) had way less occurrences than average (25% less than average). All the distributions of SHAP values (for each session and feature) were normal (Kolmogorov-Smirnov test with pval<1e-8). As a result, SHAP values of the same experimental session resulting from evaluating different models were averaged to obtain a single value per experimental session and feature.

To evaluate the variability of the models we first trained an XGBoost model and used the train dataset to obtain the SHAP values for each feature and each experiment (of the training dataset). To check if the SHAP values were stable, we retrained the model with bootstrap resamples of the training dataset and obtained new SHAP values for the original training dataset. We used the correlation of the original SHAP values with the SHAP values resulting of bootstrapping the training data to estimate the stability of the feature. Only models which had a minimum correlation of 0.5 were used for analysis. Similarly, only models with a minimum error calculated with the .632 estimator (Efron and Tibshirani, 1997) or the mean squared error regression loss (Pedregosa et al., 2011; Virtanen et al., 2020) of 7% were used.

For features representing a measure of various direct neurons, we calculated the mean (mean), maximum (max) and minimum (min) of those measures and they were introduced in the model as different features. For features representing many neurons (as in connectivity) we only obtained the mean of those measures. 43 features were used on the models. Those features were grouped in categories (in order from Fig.2.A): for quality SNR (mean, max, min); for position: depth (mean, max, min), the Euclidean distance (without depth) between neurons (mean, max, min), the difference on depth (mean, max, min) and the distance between neurons of the ensemble E1 and E2 (mean, max, min); for variation: STD of the neuronal cursor, STD of the direct neurons recoded online (mean, max, min), STD of the direct neurons calculated offline after applying CaImAn during the whole experiment (mean, max, min) or the baseline (mean, max, min); for connectivity: the average result of Granger causality between direct neurons, same for ensemble E2 to/from ensemble E1, the percentage of those pairs that Granger causality considered possible connections (also for all direct and for ensemble E1 to/from ensemble E2), the average result of granger causality from indirect neurons to direct neurons, the percentage of those pairs that were connections and the same from direct to indirect neurons. Two other features were introduced in the model that did not belong to any category: engagement of indirect neurons to the neuronal cursor and finally a feature labelling if the session was from the IT group or the PT group. Fig.2 shows 55 features (instead of 43) after separating connectivity results for different cell-classes. Features were not separated by cell-class when introduced in the model, they were separated during analysis in measures of connectivity.

It is important to note that some features may be somehow dependent on or correlated with others. As a result, their SHAP values might get arbitrarily distributed amongst each other. However, this does not affect our analysis as our goal is not to determine the best feature for learning (a final numerical value), but to discover positive or negative contributions to learning and differences for IT and PT groups.

### Final note on selection of neurons for CaBMI control

The XGBoost/Shap approach helped us understand how to better select neurons for successful CaBMI experiments. Signal quality (SNR) was highly correlated with SHAP values (Fig.2.A). In addition, SHAP values were higher, the higher the distance among all direct neurons (Fig.2.C). However, if any 2 direct neurons were too far apart (Fig.2.D), even for neurons belonging to different ensembles (Fig.2.E), SHAP values were negative. In terms of neuronal activity, positive SHAP values arose when selecting direct neurons that were silent during the baseline acquisition but highly active during CaBMI (Fig.2.F-H). We suggest experimenters attempting CaBMI to choose direct neurons that are 50 to 100um apart from each other with high SNR and the capacity of increasing greatly their baseline activity.

## SUPPLEMENTARY FIGURES

**Supplementary Figure 1:**
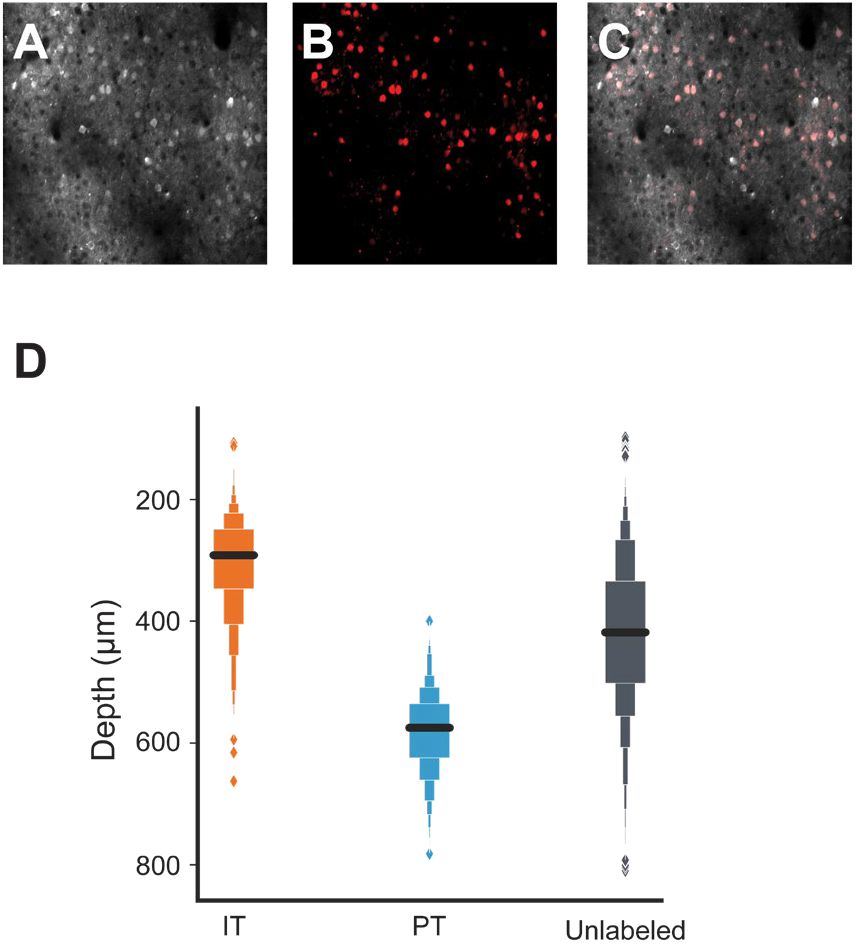
Labeling of IT and PT neurons. **A)** GCaMP6 expression under promoter Camk2a. **B)** tdTomato expressing neurons of the same plane as **A. C)** Merge of **A** and **B. D)** Boxplot of the depth of all the recorded neurons across all planes for the IT and PT group. Unlabeled neurons may belong to either cell-class in both groups.

**Supplementary Figure 2:**
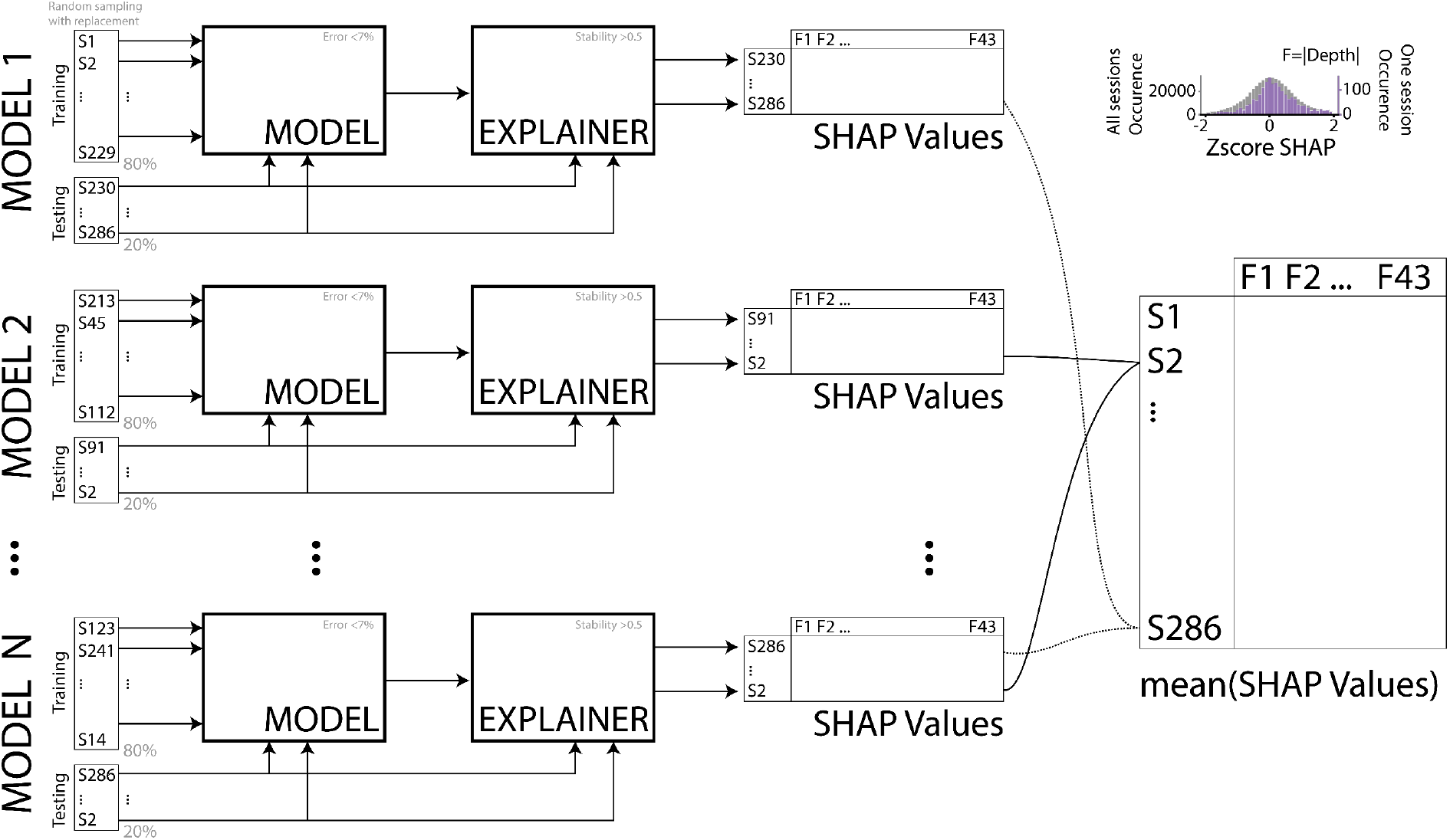
Strategy for XGBoost models and SHAP values. To obtain robust SHAP values for each session and feature, we trained XGBoost models to predict the learning readout percentage-correct for each animal and session. We only selected models (N=10000) with high accuracy and stability. Because the number of learning sessions was small relative to the number of models (286 sessions with a minimum of 15 days per animal), we trained the models with different splits of training and testing sets using random sampling with replacement. After obtaining the models, we used SHAP on each session of the testing dataset. Each of those sessions was part of a model an average of 2010 times. Thus, we averaged across all occurrences of the same session, to obtain the best approximated single SHAP value for the same session and feature. XGBoost models were calculated over all sessions jointly. SHAP values were computed on those models and separated on IT and PT sessions for some analysis a posteriori. **Top right**: Distribution of the zscore values for different occurrences of the same SHAP value across all models and sessions (grey) or all the models that included an individual example session (purple).

**Supplementary Figure 3:**
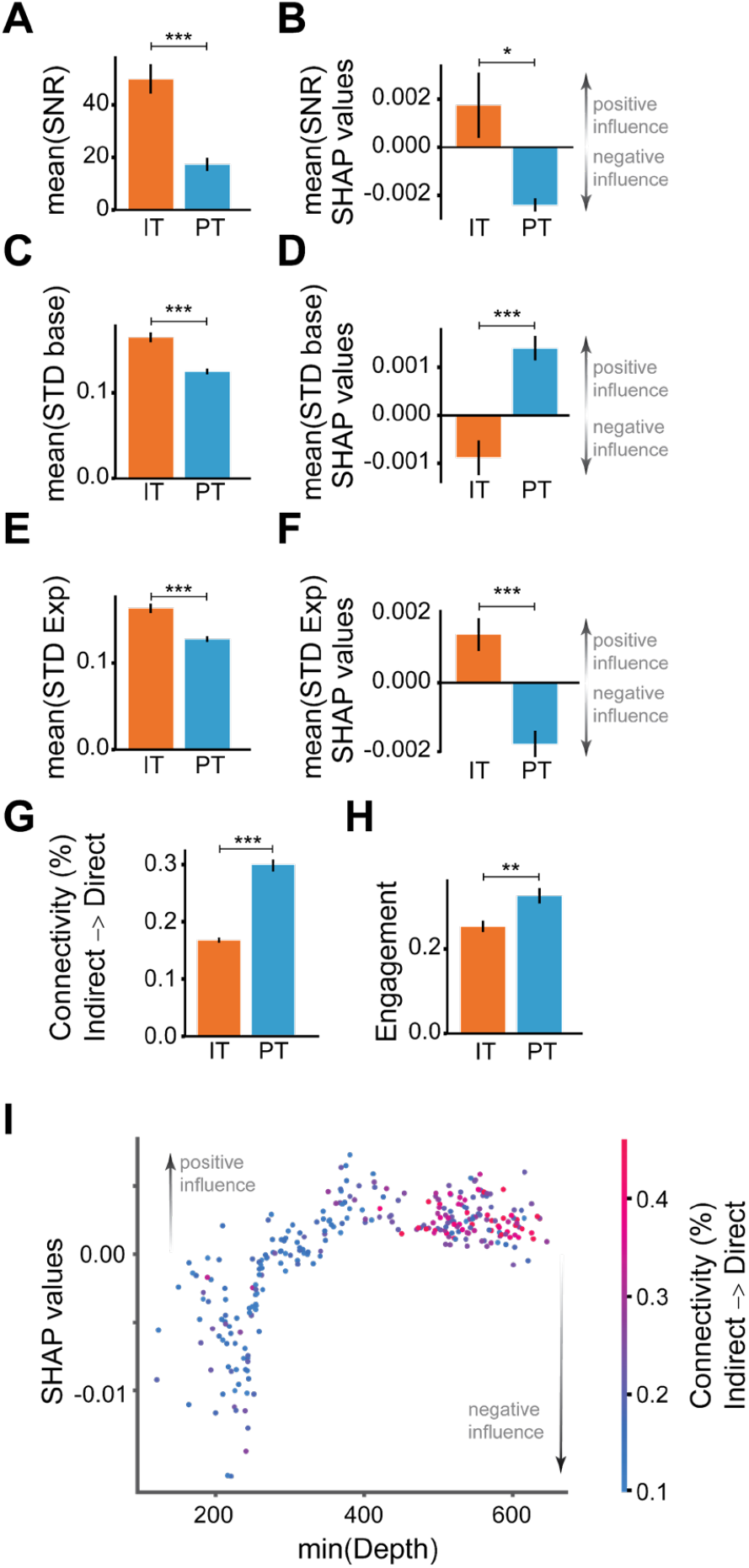
Raw value and mean SHAP values for different features. Raw value of features fed to the XGBoost model (**A, C, E, G-H**) and the mean SHAP values (**B**,**D**,**F**) of those features separated in sessions of the IT or PT groups for SNR (**A-B**); STD of the baseline (**C-D**) or the whole experiment (**E-F**). Raw value of the effective connectivity from indirect to direct neurons (**G**) and engagement of indirect neurons (**H**). IT group in orange and PT group in blue. **I)** Dependence plot between SHAP values, depth and connectivity. Colors show the value of connectivity. Each dot represents a session. Black lines in bar graphs represent SEM.(*: p<0.05,, ** : p<0.005, ***: p<0.0005 with independent t-Test).

